# Tau Oligomerization Drives Neurodegeneration via Nuclear Membrane Invagination and Lamin B Receptor Binding in Alzheimer’s disease

**DOI:** 10.1101/2025.05.21.655370

**Authors:** Nicholas Essepian, Shuo Yuan, Rebecca Roberts, Eliana Sherman, Weronika Gniadzik, Qingbo Wang, Alev Erisir, Lulu Jiang

**Affiliations:** Department of Neuroscience, University of Virginia School of Medicine, Charlottesville, VA, USA 22908; Center for Brain Immunology and Glia (BIG), University of Virginia School of Medicine, Charlottesville, VA, USA 22908; Department of Psychology, University of Virginia, Charlottesville, VA, USA

**Author notes:** These authors contributed equally (Nicholas Essepian, Shuo Yuan).

**Keywords:** Tau, Optogenetics, Nuclear Membrane, Lamina, Neurodegeneration

## Abstract

The microtubule-associated protein tau aggregates into oligomeric complexes that highly correlate with Alzheimer’s disease (AD) progression. Increasing evidence suggests that nuclear membrane disruption occurs in AD and related tauopathies, but whether this is a cause or consequence of neurodegeneration remains unclear. Using the optogenetically inducible 4R1N Tau::mCherry::Cry2Olig (optoTau) system in iPSC-derived neurons, we demonstrate that tau oligomerization triggers nuclear rupture and nuclear membrane invagination. Pathological tau accumulates at sites of invagination, inducing structural abnormalities in the nuclear envelope and piercing into the nuclear space. These findings were confirmed in the humanized P301S tau (PS19) transgenic mouse model, where nuclear envelope disruption appeared as an early-onset event preceding neurodegeneration. Further validation in post-mortem AD brain tissues revealed nuclear lamina disruption correlating with pathological tau emergence in early-stage patients. Notably, electron microscopy shows that tau-induced nuclear invagination triggers global chromatin reorganization, potentially driving aberrant gene expression and protein translation associated with AD. These findings suggest that nuclear membrane disruption is an early and possibly causative event in tau-mediated neurodegeneration, establishing a mechanistic link between tau oligomerization and nuclear stress. Further investigation into nuclear destabilization could inform clinical strategies for mitigating AD pathogenesis.

## INTRODUCTION

Alzheimer’s disease (AD) is a progressive and debilitating neurocognitive disorder characterized by pathological β-amyloid (Aβ)-containing extracellular plaques and tau-containing intracellular neurofibrillary tangles^1^, at the same time manifested by axonal withdrawal, inflammation, and accelerated neuronal degeneration ^2–4^. However, whether neurodegeneration was directly caused by these pathological hallmarks and the underlying mechanism are unclear^5^. Under normal conditions, tau promotes structural integrity along microtubules; however, in AD, hyperphosphorylation of tau leads to detachment, mislocalization, and aggregation into insoluble fibrils ^6,7^. While a significant body of literature describes the biochemical alterations of tau progression, its downstream pathophysiological effects are still largely unknown and highly debated ^8^. The neuronal protein tau is typically confined to axons through tightly regulated processes, but in tauopathies like AD and Frontotemporal Dementia (FTD), mislocalization leads to toxic aggregation and neuronal dysfunction ^8,9^. Recent studies indicate Tau oligomeric species (oTau), transitional conformations between monomers and fibrils, are the most toxic drivers of disease progression ^10–12^. These highly mobile and volatile tau species appear as an early player in AD pathogenesis and are prone to nonspecific protein interactions and prion like proliferation ^12–14^. Tau’s propensity for aberrant interactions disrupts essential cellular processes, including proteostasis and synaptic function, potentiating neuroinflammation and identifying it as a key driver of neurodegeneration ^15–17^.

An accumulating body of literature indicates that tauopathy progression is accompanied by nuclear envelope disruption ^18–21^, however exact mechanisms are still a subject of debate. Biomarkers of nuclear disturbance include tau localization to the nuclear envelope, altered protein expression patterns and defects in nucleocytoplasmic RNA transport ^16,18,22^. The nuclear envelope plays a crucial role in maintaining genomic stability by regulating chromatin organization, transcriptional activity, and nucleocytoplasmic trafficking ^23^, so disruptions can lead to widespread alterations in gene expression, potentially exacerbating neurodegenerative processes ^24^. Furthermore, recent work shows pathogenic tau to be associated with lamina destabilization including nuclear invaginations and blebbing ^18,25,26^.

Latest work from our lab indicates that oligomeric Tau binds directly to LaminB2 and LaminB receptor proteins, likely rendering the proteins insoluble and delocalized ^20^. The nuclear lamina in nonpathological neurons is crucial to inner nuclear envelope structure and assists in chromatin organization, DNA replication, and mediation of the nucleocytoplasmic interface ^27–29^. Curiously, aggregate disruption of the lamina nucleoskeleton is not confined to AD pathology, as similar findings are demonstrated in Huntington’s disease (huntingtin protein) ^30^ and FTD where tau aggregates are shown to recruit microtubules to nuclear malformations ^25^. AD studies have demonstrated that induction of pathogenic tau decreases nuclear tension ^21^, induces structural changes in the LaminB nucleoskeleton ^18,26^ and recruits LaminB components to tau oligomers ^19,20^. Therefore, it is likely that pathological tau aggregates invade into the nuclear envelope where they bind to and delocalize nuclear lamin components, thereby inducing nuclear malformations and chromatin reorganization.

To address the question of how tauopathy affects nuclear disruption and neuron death in AD, we track the dynamics of tau aggregation in human iPSC derived cortical neurons as well as in the brains of humanized P301S tau transgenic (PS19) mice during disease progression. In neuronal cell culture, we used the genetically engineered construct 4R1N Tau::mCherry::Cry2Olig system to induce tau oligomerization under exposure of 488λ blue light ^19^. We find that tau granules under active oligomerization aggregate towards the nuclear envelope and induce significant intrusions into the nuclear space. These findings are reflected in mouse brains from postnatal month 5 (PM5) and postnatal month 9 (PM9) PS19 mice where pathological tau accumulation within the nuclear bounds strongly correlates with degree of lamina disruption. Electron microscopy finds progressive chromatin decompaction under tauopathy conditions in PS19 brains. Notably, our findings from the in vitro cell culture and in vivo mouse models are validated in human postmortem brain tissue from AD patients spanning Braak stages 1-6. Collectively, our findings demonstrate tau oligomer nucleo-toxic effects proceeding and potentiating neuronal dysfunction seen in AD.

## RESULTS

### Live cell imaging of iPSC induced neurons reveals OptoTau oligomer granules localize to the nuclear envelope and induce invagination under 488λ blue light exposure

To examine the effects of tau oligomerization in real time, we used iPSC induced neurons transduced with either 4R1N Tau::mCherry::Cry2Olig (OptoTau) or mCherry::Cry2Olig (mCherry) constructs ^19^. These systems use fluorescent mCherry linked with the bacterial protein Cry2Olig which dimerizes under 488λ blue light exposure (**Fig. 1A**), however, the mCherry control vector has no additional linked protein while OptoTau expresses the 4R1N tau isoform, a key mediator of AD tau toxicity ^19,31^. This means that while mCherry forms temporary dimers of Cry2Olig, OptoTau facilitates prolonged stress granule formation by facilitating stable oligomerization of 4R1N tau proteins (**Fig. 1A**). This makes OptoTau a powerful tool to induce and observe tau stress granules in a controlled time course (**Fig. 1B**). In addition to mCherry fluorescence, we used the cell membrane permeable DNA dye Nucspot Live 488 designed for live cell imaging applications to mark the nuclei.

**Fig. 1.**
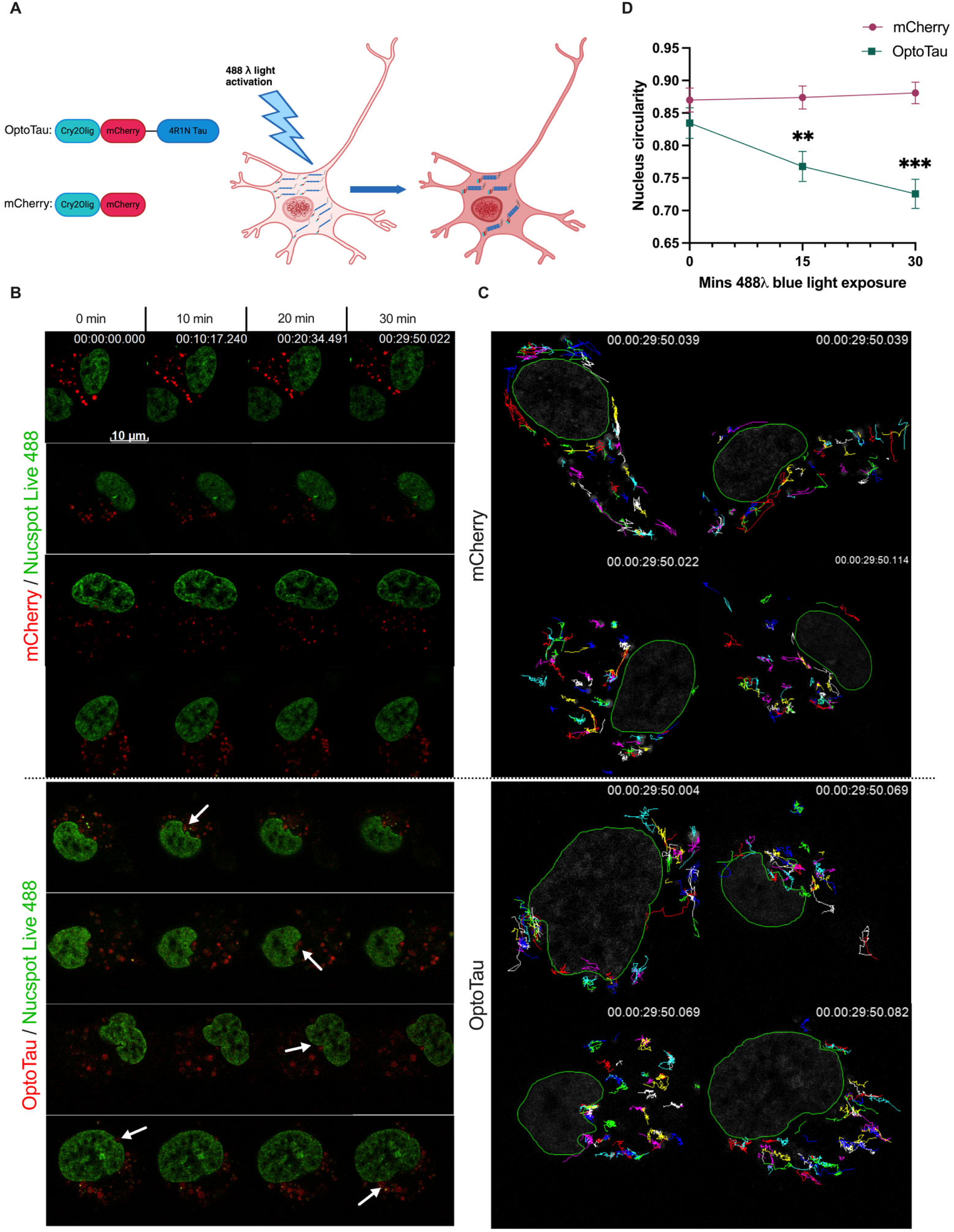
Live cell confocal imaging of iPSC derived neurons demonstrates OptoTau aggregate localization to the nuclear envelope followed by envelope misshaping. **A.** Schematic illustrating of OptoTau and mCherry live cell activation vector transcripts followed by mechanism of OptoTau oligomer formation under 488λ blue light exposure. **B.** Representative timelapse images of mCherry and OptoTau expressing neurons stained with Nucspot Live 488 at 0, 10, 20, and 30 minutes of blue light exposure. **C.** Closeup representative images of neuronal nuclei at 30-minute timepoint overlaid with particle tracking lines following mCherry or OptoTau fluorescent granules over time course. Green outline denotes the nucleus at 30 minutes. **D.** Mean nucleus circularity at 0, 15, and 30 minutes of blue light exposure. *n* = 9-10. ** = *p* < 0.01, *** = *p* < 0.001. Error bars indicate 95% standard error of the mean.

Previous studies of AD and similar neurodegenerative conditions have found atypical protein aggregates to cause nuclear envelope abnormalities including destabilization, invagination, leakiness, and rupture ^16,20,21,25,30,32^. Consistent with established findings, 30 minute blue light time courses exhibit OptoTau aggregate localization to the nucleus and nuclear misshaping (**Fig. 1B, Video S1**). Neurons with mCherry control show presence and movement of fluorescent granules within the cell body but granular motility was largely random and did not localize at the nucleus (**Fig. 1B**). Particle tracking over 30 minutes confirms mCherry granules exhibit random movement patterns within cell bodies and around the nuclear envelope while OptoTau aggregate progressive lines frequently converge toward forming cavities in the nucleus and in some cases cross the nuclear boundary (**Fig. 1C**). Further analysis at 0, 15, and 30 minute time points reveals that although mCherry and OptoTau nucleus circularity begins at similar means of 0.87 and 0.83, respectively, OptoTau nuclear circularity decreases significantly with blue light exposure while mCherry remains stable (**Fig 1D**). This finding aligns with the observation that fluorescent OptoTau aggregates induce concavities in the nuclear envelope during blue light activation (**Fig. 1B**). More importantly, this result confirms that nuclear abnormalities are due to active oligomerization of 4R1N tau and not mCherry::Cry2Olig vehicle proteins.

### iPSC induced neurons expressing OptoTau show multiple elevated pathological tau biomarkers following prolonged blue light activation

Previous work done by our lab has established proper functionality of OptoTau oligomerization in mouse derived primary cortical neurons ^19^, but the use of iPSC derived neurons offers a physiologically relevant platform for studying human AD ^33,34^. iPSC derived neurons have the advantage of developing human neuronal features over time, including synapse formation, cell morphology, and response to neurodegenerative stressors such as tau accumulation ^33,34^. To assess OptoTau efficacy in studying AD tau pathology in vitro, we exposed transduced neurons expressing OptoTau or mCherry with 0 or 60 minutes of 488λ blue light and probed for various pathological tau biomarkers in fixed cells. Additionally, all cells were IHC stained for dsDNA using Dapi, mCherry to confirm vector expression, and TUJ1 to confirm neuronal identity (**Fig. 2A-D**).

**Fig. 2.**
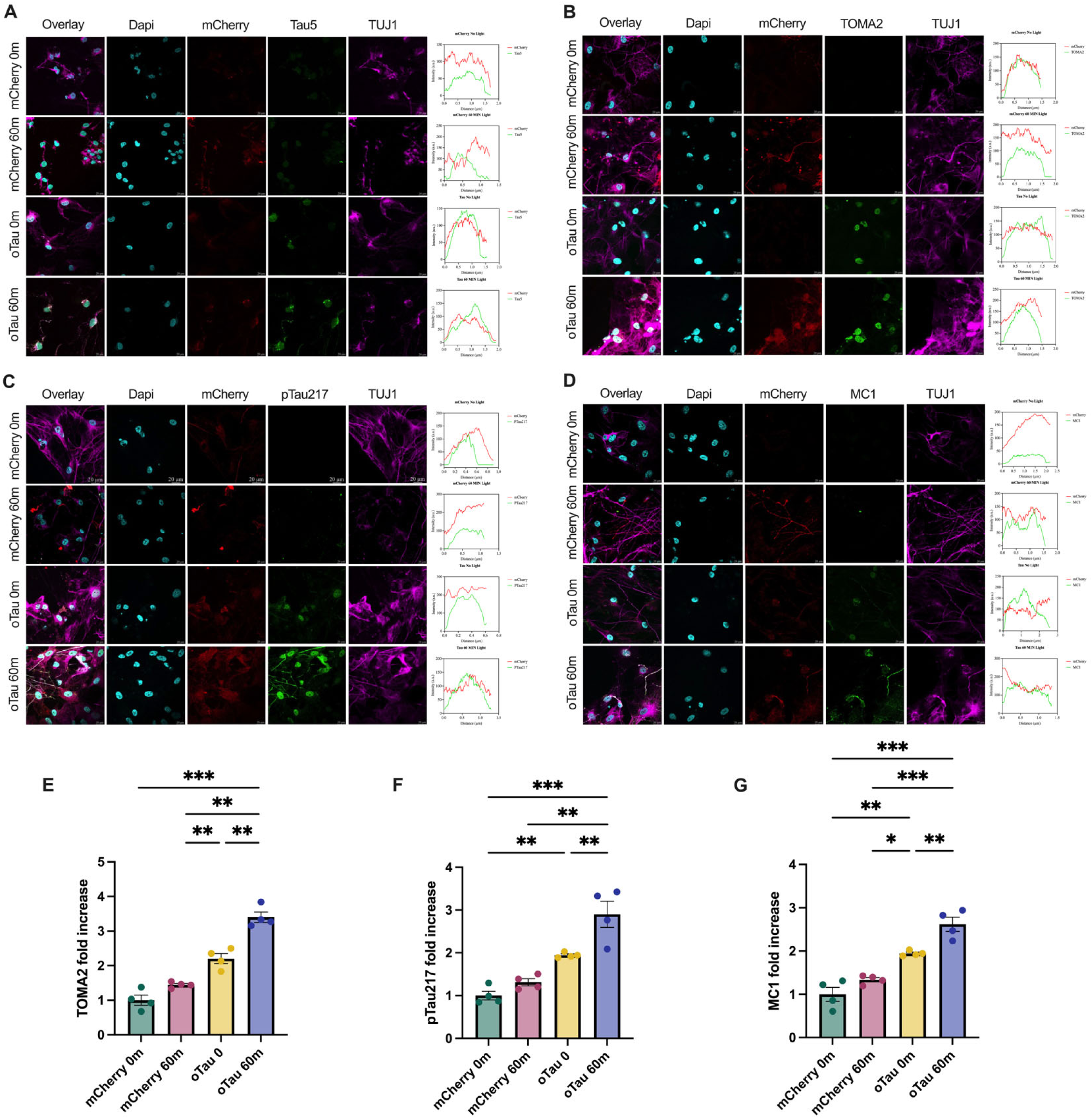
OptoTau expressing neurons show pathological tau accumulation following prolonged 488λ blue light activation examined by multiple tau markers. **A.** Representative confocal images of fixed mCherry and OptoTau induced neurons following 0 or 60 min 488λ blue light activation at 63x magnification. IHC staining marks dsDNA stain Dapi, construct marker mCherry, global tau marker Tau5, and beta tubulin neuronal marker TUJ1. Representative histograms show fluorescence intensity of mCherry and tau marker colocalization in a single cell. **B.** IHC staining and representative histograms using oligomeric tau marker TOMA2. **C.** IHC staining and representative histograms using phosphorylated tau marker pTau217. **D.** IHC staining and representative histograms using misfolded tau marker MC1. **E.** TOMA2 oligomeric tau fluorescence intensity fold increase in representative ROIs. **F.** pTau217 phosphorylated tau fluorescence intensity fold increase in representative ROIs. **G.** MC1 misfolded tau fluorescence intensity fold increase in representative ROIs. Error bars indicate 95% standard error of the mean. * = *p* < 0.05, ** = *p* < 0.01, *** = *p* < 0.001. Analysis was conducted using one-way ANOVA with Tukey’s multiple comparisons.

To assess overall tau accumulation, we stained using Tau5 that detects global tau ^35^. Tau5 fluorescence showed strong increases in OptoTau neurons with 0 or 60 minutes of activation and strong single cell colocalization when compared with mCherry control neurons (**Fig. 2A**). This confirms total tau enhancement under oTau transduction and indicates reliable expression of both constructs. Next, we assessed oligomeric tau using TOMA2 markers specifically targeted for neurotoxic tau aggregates in AD ^36^. As expected, TOMA2 intensity appeared highest in light activated OptoTau neurons and showed moderate histogram colocalization (**Fig 2B**). Fluorescence quantification revealed greatest fold increases in TOMA2 among exposed OptoTau (**Fig. 2E**), confirming greater tau oligomer formation.

Further probing with pTau217, a marker for tau phosphorylation at threonine 217 which correlates closely with AD progression ^37^, showed close fluorescence peak overlap in OptoTau neurons following activation (**Fig. 2A**). pTau217 fluorescence also showed highest levels in activated oTau neurons (**Fig. 2D**). Phosphorylation at Tau-217 is increasingly found to be an early and specific modification in AD and is present in both detached monomers oligomers ^37,38^. This, in combination with its high colocalization with oTau granules makes it a suitable biomarker for tracking accumulation. Finally, misfolded tau species were assessed using the highly specific MC1 antibody that recognizes a pathological tau conformation only seen in AD afflicted neurons ^39^. Once again, highest colocalization histograms and fluorescence fold increases presented in OptoTau neurons after 60 min of blue light exposure. Across all markers, we see higher colocalization between mCherry and pathological tau signals as well as significant increases in tau among treatment cells, suggesting OptoTau activation drives the formation of disease relevant tau oligomers, phosphorylation, and conformations. Together, these results indicate that light induced OptoTau activation showed promising AD tauopathy recapitulation in iPSC neurons.

### iPSC induced neurons show higher incidence of nuclear disruption under tau oligomerization

Recent studies indicate that accumulation of pathological tau around the nucleus can induce physical distortions including nuclear invaginations, nuclear pore complex (NPC) impairment, and lamina dysfunction ^18,20,21,25^. These findings are increasingly demonstrated across neurodegenerative tauopathy models including AD, and FTD ^20,25^. Similar findings of lamina invagination in non tau related conditions such as Huntington’s disease, involving the polyglutamine aggregates ^30^, increasingly suggest an overarching category of laminopathies – neurodegenerative conditions presenting with increasing lamina layer disfunction. Given the role of lamina proteins, particularly Lamin B2 (LB2) and Lamin B receptor (LBR), in directing chromatin architecture in neurons, elucidating patterns of nuclear disruption is critical in understanding molecular bases of disease progression in AD and related laminopathies^40–42^. Based on our observations in livecell timelapse we utilized the same mCherry and OptoTau constructs to examine disruption in fixed iPSC induced neurons. Neurons transduced with mCherry and OptoTau were exposed to 0 or 60 minutes of 488λ blue light and IHC stained for DNA, lamina components, tau isoforms and neuronal markers (**Fig. 3A**).

**Fig. 3.**
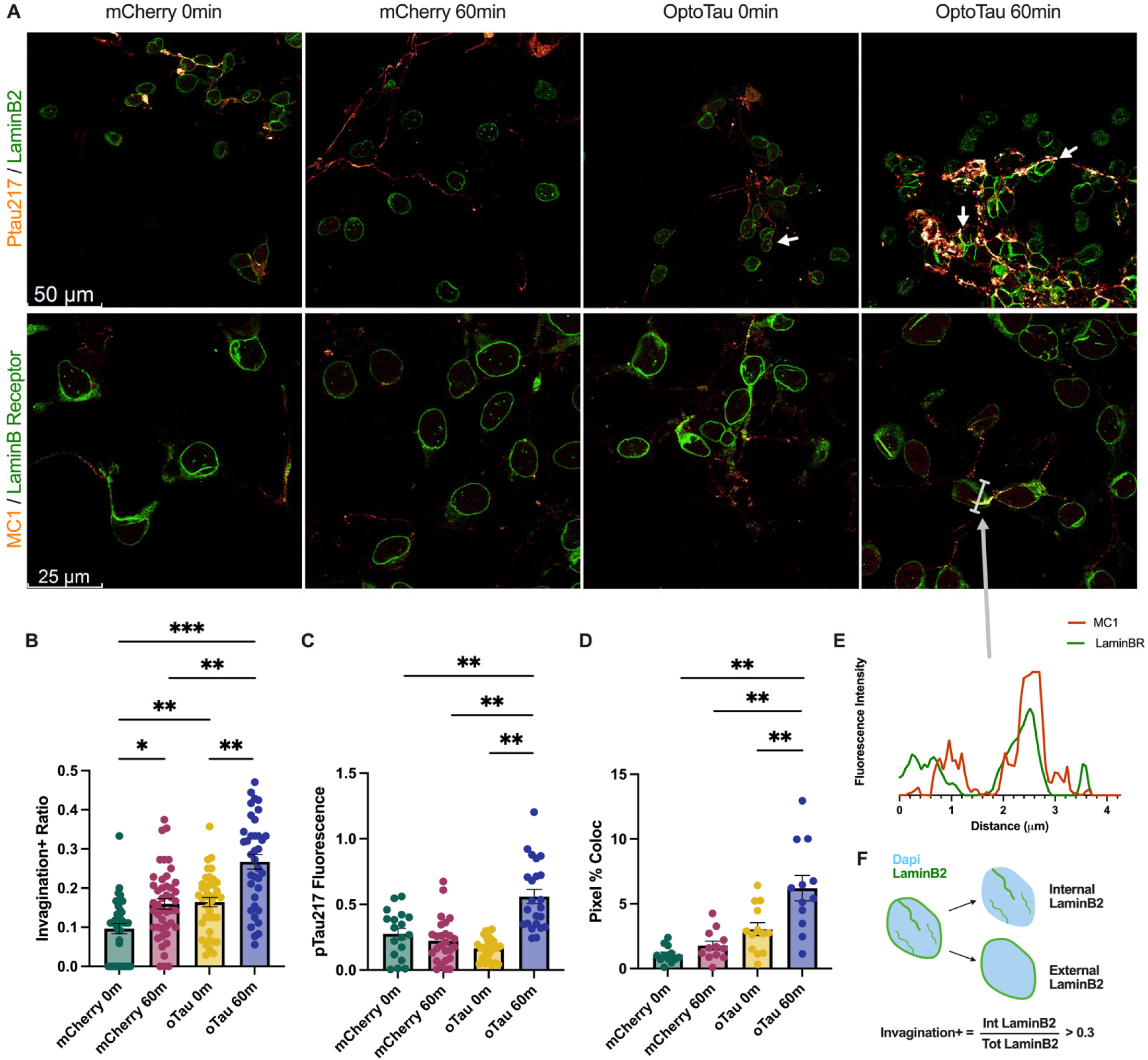
Fixed cell imaging finds higher prevalence of nuclear envelope invagination and greater lamina/tau colocalization in OptoTau expressing neurons following prolonged 488λ blue light exposure. **A.** Representative confocal images of fixed mCherry and OptoTau induced neurons at 63x (top) and 63×2 (bottom) magnifications. **B.** Ratio of nuclei that present with invagination within each ROI containing 10-60 cells. Dataset derived from two separate rounds of fixing, staining, and imaging; *n* = 37-45. **C.** Fluorescence intensity of pTau217. Higher fluorescence corresponds to integrated density normalized to Dapi dsDNA staining. *n* =18-26. **D.** Colocalization analysis of pixel % overlap between pTau217 and LaminB2; *n* = 12-13. **E.** Fluorescence intensity histogram of nuclear envelope invagination example along bar indicated in representative image. X axis in units of microns (μm). **F.** Schematic illustrating criteria for invagination: nuclei with internal (overlapping with dapi staining) LaminB2 or LBR exceeding a proportion of 0.3 of total LaminB2or LBR are counted as invaginated. Error bars indicate 95% standard error of the mean. * = *p* < 0.05, ** = *p* < 0.01, *** = *p* < 0.001. Analysis done using one-way ANOVA with Tukey’s multiple comparisons.

Our results indicate commonalities between live cell observations and fixed cell nuclear morphologies. Using fluorescence mapping of dsDNA stain dapi and lamina proteins, LB2 and LBR, nuclei were evaluated for presence of lamina folds overlapping with internal nuclei, indicative of nuclear invagination. Nuclei with internal LB2 at a ratio greater than 0.3 of total LB2 at each nucleus qualified as invaginated (**Fig. 3F**) ^25^. Examining ratios of nuclei presenting with invagination in each ROI, we find highest rates of nuclear invagination in OptoTau cell culture following prolonged blue light activation (**Fig 3B**). This finding indicates that upon oligomerization, tau granules localize to the nuclear membrane and exert mechanical pressure on the lamina nucleoskeleton, potentially impairing integrity. These results were validated with LB2 and LBR staining, indicating mislocalization of critical nuclear envelope proteins. Furthermore, activated OptoTau neurons presented with elevated levels of phosphorylated tau, aligning with established mechanisms of tau oligomer proliferation by post translational modification (**Fig. 3C**)^37,38^.

As further evidence for interaction between tau and lamina components, OptoTau neurons following activation showed elevated levels of colocalization between phosphorylated tau and LaminB2 (**Fig. 3D**). Fluorescence intensity mapping of one example of invagination shows direct overlap of MC1 misfolded tau marker with LBR invagination (**Fig. 3E**). These findings support our proposed hypothesis of lamina disruption by tau oligomer localization to the nuclear envelope. Presence of lamina aberrations and nuclear strain in neurons has the potential to drive neurodegeneration via nuclear stress responses, disrupted chromatin structure, protein expression profiles, and nucleocytoplasmic transport.

### Mouse neurons under tau conditions present higher prevalence of nuclear invaginations

To examine LaminB2 disruption in tau mouse conditions, we utilized C57 BL/6 wild type (WT) and P301S tau (PS19) transgenic mouse strains as baseline and tau pathology mouse models, respectively. The PS19 mouse model harbors the P301S human mutation to MAPT with one N terminal insert and four microtubule binding repeats (4R1N) ^43–45^. PS19 mice are a well-established transgenic mouse strain presenting mature tau pathology and recapitulating AD tauopathy progression ^43,45,46^. Utilizing a robust mouse model allows for the comparison of cell culture results with an in vivo brain application over the course of disease progression. As such, we harvested brain slices from PM5 and PM9 WT and PS19 mice and probed for similar target proteins as transduced cell culture neurons (**Fig. 4A**)

**Fig. 4.**
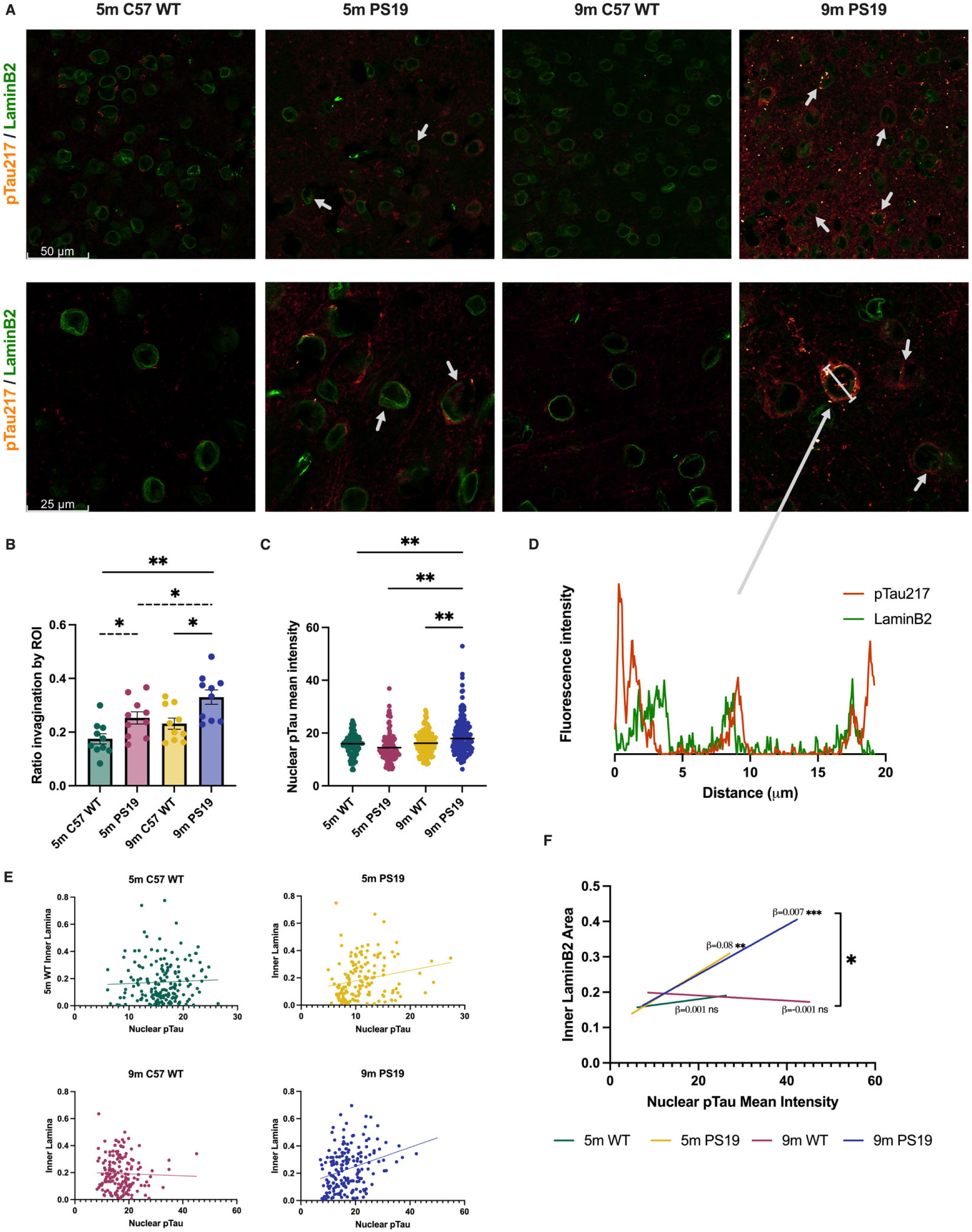
Tau mouse neurons present higher prevalence of nuclear lamina invagination. **A.** Representative confocal images of postnatal month 5 (PM5) and postnatal month 9 (PM9) neurons from C57 WT and PS19 tau frontal cortex, 63x (top) and 63×2 (bottom) magnifications. **B.** Ratio of nuclei that present with invagination (internal LaminB2 ratio ≥ 0.3). **C.** pTau217 mean fluorescence intensity within the nuclei of PM5 and PM9 neurons. **D.** Representative fluorescence intensity histogram of nuclear envelope invagination example along bar indicated in representative image. **E.** Correlation between mean pTau intensity within the nuclei and inner lamina proportion. **F.** Correlation slopes from Fig. c plotted on a combined axis. *n* = 158 to 177 nuclei. For significantly nonzero slope of 5m WT, 9m WT, 5m PS19, 9m PS19, *p* = 0.56, 0.68, 0.0078 (**), 0.0001 (***) respectively. For, difference in slopes, *p* = 0.012. Error bars indicate 95% SEM. * = *p* < 0.05, ** = *p* < 0.01. Solid lines indicate one-way ANOVA; dashed indicate unpaired *t* test.

Our results indicate findings consistent with neuronal cell culture. 5 and 9 month PS19 neurons show higher levels of phosphorylated tau than their WT counterparts (**Fig 4A**). Mouse neurons also present notable incidence of misshapen and invaginated nuclei with localization of tau (**Fig. 4A**). As in Fig. 3, nuclei were evaluated for invagination by quantification of LaminB2 inside the nucleus, indicative of nuclear folding. Nuclei with internal LaminB2 at a ratio of 0.3 or more of total LaminB2 qualify as invaginated. Quantification of ROI in WT and PS19 cortex reveals significant increases in incidence of nuclear invagination among 5 and 9 month PS19 mice over age matched controls (**Fig. 4B**). Notably, rates of nuclear disruption are comparable between 5m PS19 and 9m WT, indicating a trend of accelerated neuron aging through lamina disruption^47,48^.

Mapping pTau217 fluorescence within individual nuclei using similar quantification methods, we find that mean intensities of phosphorylated tau were highest among 9m PS19 as well (**Fig. 4C**). As observed in Fig. 5a, visible folds within the lamina layer show incidence of tau localization to nuclear folds as observed in cell culture. A fluorescence intensity histogram of one example reveals high intensity pTau in close proximity to outer lamina and direct overlap of pTau with LaminB2 in an internal nuclear membrane fold (**Fig. 4D**). Finally, correlating nuclear pTau217 mean intensity with internal LaminB2 of individual nuclei reveals that only 5 and 9 month PS19 mice show significantly nonzero correlations (*p* = 0.0078, 0.0001 respectively) and are significantly greater than their WT counterparts (**Fig. 4E, F**). Taken together, these findings align with our hypothesized mechanism of pathological tau directly binding and disrupting nuclear lamina components. More importantly, it demonstrates that disruptions found in tau-induced cell culture replicate in live animal brains under tauopathy conditions.

**Fig. 5.**
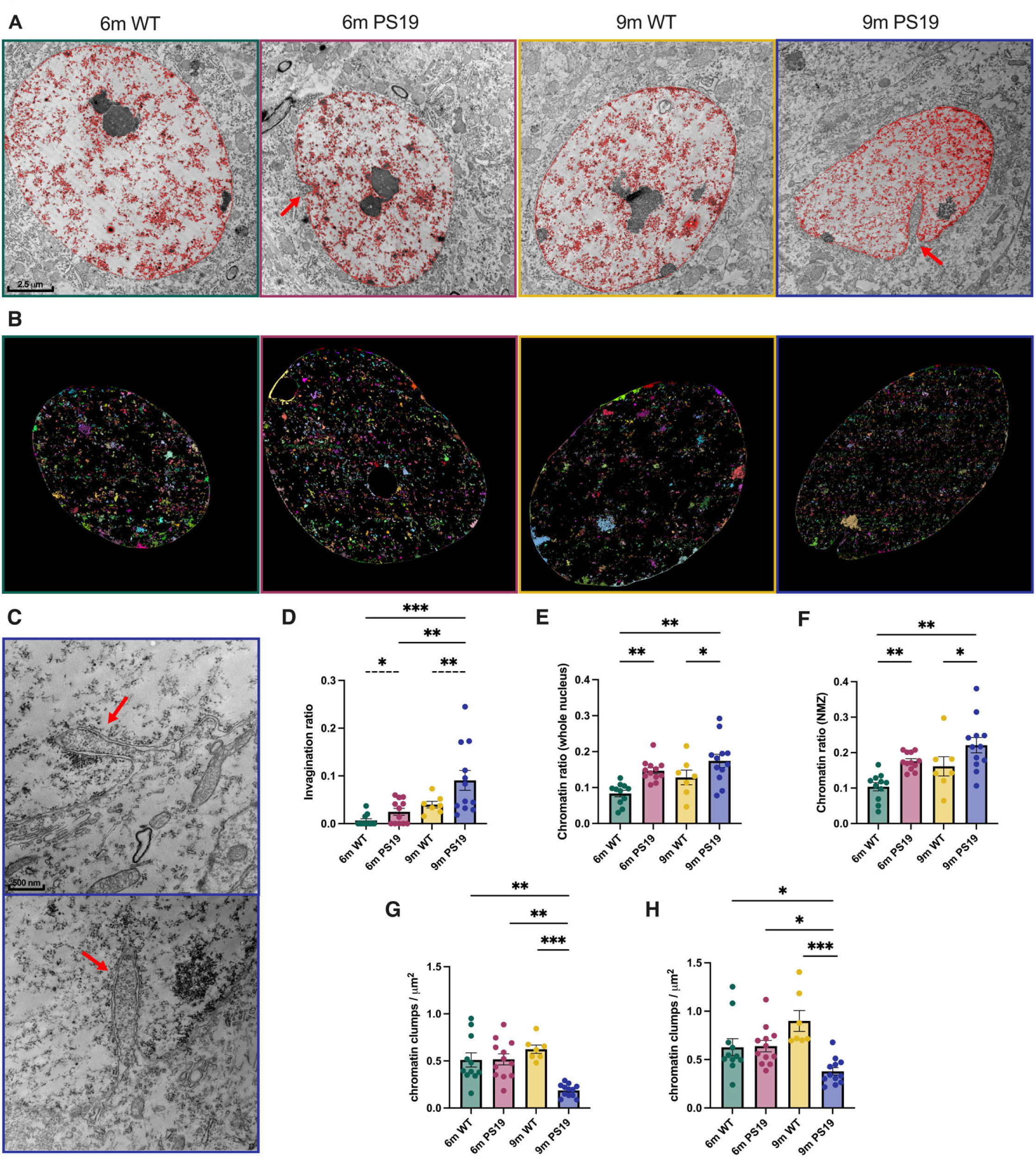
Hippocampal neurons of PS19 mouse brain with accumulated tau pathology demonstrate changes in nuclear chromatin density. **A.** Representative electron microscopy (EM) images of PM6 and PM9 WT and PS19 CA1 neurons at 4kx magnification. Chromatin outlined in red. **B.** Representative connected components labeling of chromatin at 4kx magnifications. Components are defined as closely connected pixels. **C.** Representative EM images of PM9 nuclear invaginations at 15kx magnification. **D.** Ratio of invagination nuclear membrane length relative to the perimeter of the whole nucleus. **E.** Chromatin area coverage relative to the entire nucleus excluding nucleoli across conditions. **F.** Chromatin area coverage relative to the area of the nuclear membrane zone (NMZ), the outermost 20% region of the nucleus along the inside of the nuclear membrane. **G.** Density of chromatin clumps normalized to whole nucleus area excluding the nucleoli. Clumps are chromatin components with an area ≥0.04 μm^2^. Lower values correspond with lower chromatin clump density. **H.** Density of chromatin clumps within the NMZ normalized to the NMZ area. Error bars indicate 95% SEM. *n* = 8 to 14. * = *p* < 0.05, ** = *p* < 0.01, *** = *p* < 0.001. Analysis was done using unpaired t-tests, one-way and 2-way ANOVA with Tukey’s multiple comparisons.

### Mouse neurons under tau conditions show severe nuclear morphological anomalies and evidence of chromatin reorganization

Building on our prior results finding elevated rates of nuclear invagination in PS19 tau mouse brains over age matched controls (**Fig. 4**), we used transmission electron microscopy (EM) to examine ultrastructural abnormalities in both nuclear shape and chromatin density of PM6 and PM9 C57 WT and Tau P301S PS19 mice at high magnifications of 4kx and 15kx. Representative EM images of cortical neuron nuclei reveal emergence of nuclear envelope abnormalities including large invaginations protruding into to the intranuclear space in PS19 tau mouse brains (**Fig. 5A, C**). Quantification finds invaginations to be of the greatest incidence and relative size by membrane length in 9-month PS19 mice, however smaller intrusions also appear in 6-month PS19 mice. (**Fig. 5D**). Notably, invaginations similar in incidence and size to 6-month PS19 mice also appeared in 9-month WT mice, in line with previous reports of nuclear lamina folding as part of physiological aging^47,48^. These studies found Lamin B1 invaginations in neurons to be a part of natural CNS aging ^47^, however, the more frequent and exaggerated invaginations in age matched tau mice suggests a disease-accelerated aging phenotype.

Given the importance of lamina stability in maintaining genomic integrity during aging ^47–49^, we investigated whether lamina structural aberrations coincide with broad changes in chromatin organization. Previous studies have indicated that tauopathy progression is linked with genomic reorganization, particularly chromatin relaxation marked by decreased heterochromatin content ^50,51^. Using chromatin area coverage and connected components labeling analysis, we assessed chromatin distribution within the nuclei. Consistent with established findings, PS19 neurons from 6 and 9-month mice showed global chromatin loosening compared with age matched controls (**Fig. 5E, G**). This was reflected in increased chromatin coverage across the nucleus (**Fig. 5E**) and decreased chromatin clump density in 9-month tau mice, indicating tauopathy promotes heterochromatin loss and nuclear decompaction, particularly in neurons presenting with greatest lamina disruption (**Fig. 5G**). To further localize the effects of LaminB2 disruption, we defined a nuclear membrane zone (NMZ) as the outermost 20% of the nucleus to capture changes in perinuclear chromatin rich is lamina associated domains (LADs) ^47,52^. Within the NMZ, PS19 neurons exhibited increased chromatin coverage compared with age matched controls (**Fig. 5F**). The NMZ also showed greater relative decreases in chromatin density in the outer nucleus (**Fig. 5H**), further indicating increased euchromatin dispersal in response to nuclear invagination related stress. Since this zone is highly enriched in LADs, it would make sense that lamina destabilization would impair genome anchoring.

Together, these findings provide confirmation that tau pathology induces ultrastructural changes in nuclear envelope morphology, accompanied by global chromatin reorganization. Decreased heterochromatin content, particularly in the NMZ, further implicates lamina dysfunction in the degradation of genome organization. Altered genetic architecture may underlie aberrant transcription patterns observed in AD that mediate accelerated degeneration ^51,53,54^. Lastly, this data reinforces the idea of tau-driven neuronal aging acceleration and highlight chromatin as a key target of tau toxicity.

### Disrupted nuclear integrity of Lamin B receptor correlates with early misfolded tau emergence in human Alzheimer’s disease brain tissue

To ensure our findings in cell culture and tau mouse models show consistency in AD progression, we examined human postmortem brain tissue from AD patients using IHC single staining with DAB. We probed for misfolded tau using MC1 ^39^ and Lamin B receptor (LBR) as we had previously done in cell culture to track pathological tau accumulation and nuclear disruption. Representative images from Brodmann area 10 of the prefrontal cortex show early but mild presentation of MC1 tau beginning in Braak stages 3-4, which in this dataset are still clinical controls, and markedly increased tangle accumulation in stages 5-6 (**Fig. 6A**, top panel). Quantification of MC1 by DAB intensity confirms sharp increases among stages 5 and 6 compared with stages 1-4 (**Fig. 6B**). Further analysis by clinical controls and AD further confirms significantly elevated MC1 misfolded tau pathology in late-stage AD patients (**Fig. 6C**), however it should be noted that small quantities of tau aggregates are visible in stages 3 and 4 (**Fig. 6A**). These results confirm well established findings of smaller tau oligomers early in disease progression with large tau complexes presenting in late stage AD pathology ^55,56^.

**Fig. 6.**
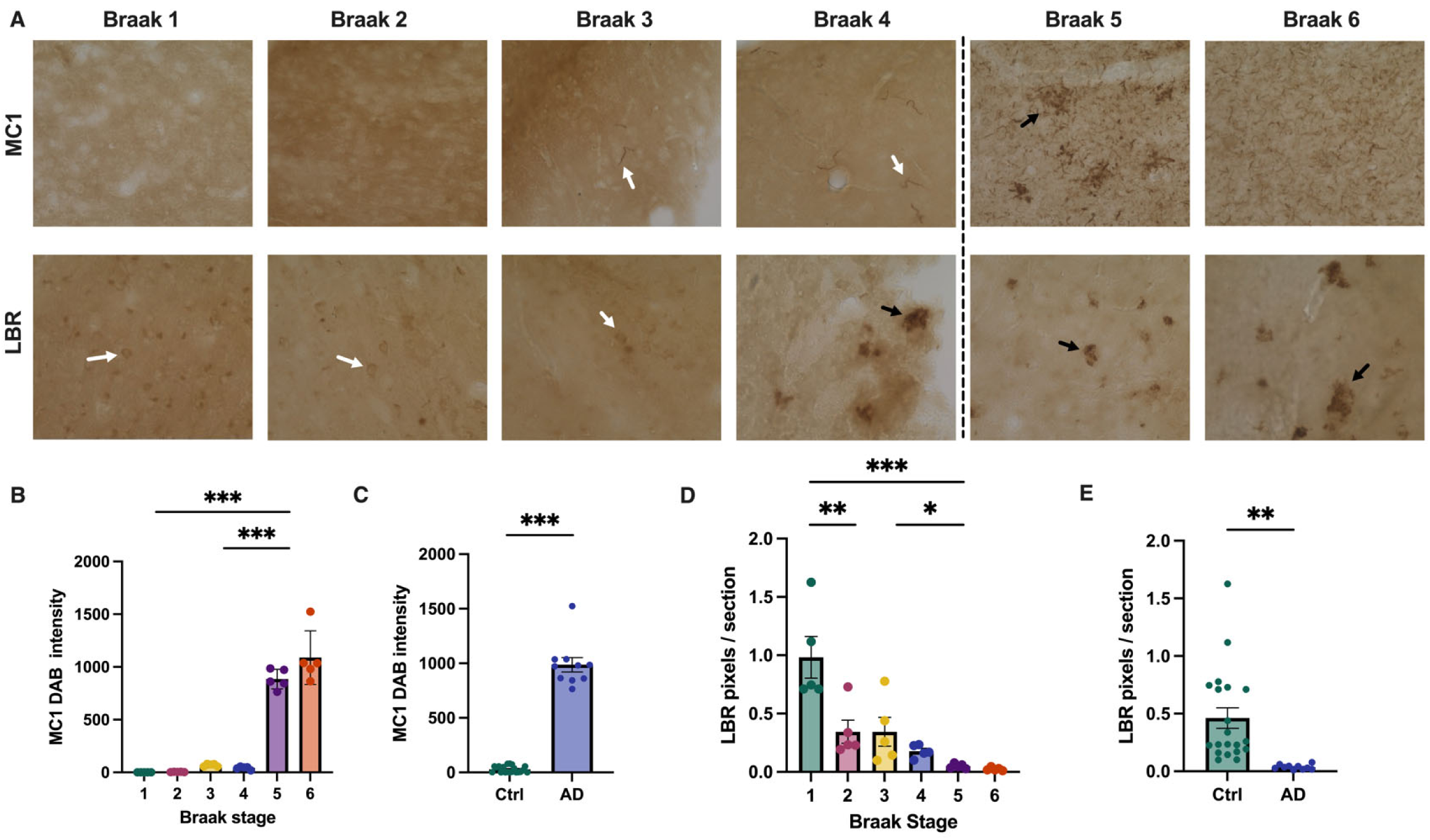
IHC staining of human AD postmortem brain tissue presents disrupted Lamin B receptor nuclear integrity in early AD pathogenesis. **A.** Representative images of human AD postmortem brain tissue, Braak stages 1-6, taken at 40x magnification. Tissue stained by IHC with DAB for MC1 (top) and Lamin B receptor (bottom). Stages 1-4 in this data set were clinical controls while 5 and 6 are advanced stage AD. **B.** Normalized integrated density of MC1 (misfolded tau) IHC staining with DAB by Braak stage. *n* = 5. **C.** Normalized integrated density of MC1 (misfolded tau) staining by control vs disease, *n* = 20, 10. **D.** Nuclear integrity of Lamin B receptor using IHC staining with DAB by Braak stage, *n* = 5. Integrity is calculated as circular continuity of pixels within each ROI. **E.** Nuclear integrity of Lamin B receptor staining by control vs disease, *n* = 20, 10. Error bars indicate 95% SEM. * = *p* < 0.05, ** = *p* < 0.01, *** = *p* < 0.001. Analysis was conducted using unpaired t tests and one-way ANOVA with Tukey’s multiple comparisons.

Strikingly, LBR staining demonstrates progressive loss of nuclear integrity in tandem with early misfolded tau emergence (**Fig. 6A**, bottom panel). In early control Braak stages 1-3, LBR shows high nuclear circularity, indicative of intact nuclear envelopes, whereas stage 4 tissue shows the emergence of disrupted, fragmented staining patterns, reflective of compromised nuclear morphology (**Fig. 6A**, bottom panel). Quantification of nuclear integrity calculated as circular continuity within nuclear ROIs reveals significant decreases in LBR nuclear integrity as early as Braak stages 2-3, with continued decline into later stages (**Fig. 6D**). Analysis by clinical status confirms significantly decreased nuclear LBR integrity from control to AD brain tissue (**Fig. 6E**). Notably, quantified disruption emerged concurrently with mild MC1 tau aggregation and became compromised prior to severe fibril accumulation. These findings suggest that nuclear envelope disruption occurs in early AD pathogenesis and demonstrates consistency with nuclear morphology differences observed in cell culture and AD mouse models. Initiation of nuclear disturbance by tau oligomers prior to extensive fibril accumulation supports our hypothesis that lamina interference may be an initiating factor in neuronal dysfunction. Furthermore, our observations align with prior findings of lamina disruption in AD and related neurodegenerative tauopathies ^18,20,25^. This, in combination with the consistency between human tissue, cell culture neurons, and AD tau mouse models further reinforces the role of nuclear destabilization as a pathogenic feature of Alzheimer’s disease.

## DISCUSSION

While the role of tau aggregation has been extensively studied in its effects on neurodegeneration, the precise mechanisms by which tau oligomers induce cellular dysfunction and toxicity remains poorly understood. Furthermore, the role of nuclear destabilization, a process observed in AD and similar tauopathies, in initiating and driving disease progression is uncertain. The findings presented in this study provide compelling evidence that tau oligomer species initiate and drive nuclear envelope disruption as an early and contributing mechanism in Alzheimer’s disease pathogenesis. Utilizing a robust, multimodal approach with human iPSC induced neurons in cell culture, transgenic mouse models, and human AD postmortem brain tissue, we find that tau oligomers localize to and interact with nuclear lamina proteins Lamin B2 and Lamin B receptor, thereby leading to invagination, compromised nuclear integrity, and chromatin decompaction. These findings align with recent studies implicating tau in toxic effects on the nucleus ^18–20,25^ and offers new insight into early pathological mechanisms of aggregate-lamina disruption across multiple neurodegenerative diseases.

The use of optogenetically activated 4R1N tau vectors in iPSC derived neurons displays that tau oligomerization, an early process in tauopathies ^14,37,38,56^, leads to rapid and significant nuclear envelope deformation under blue light induction. The decreases in nuclear circularity observed in 30 minutes of activation were specific to OptoTau expressing neurons, confirming resulting structural abnormalities are not artifacts of Cry2Olig activation or protein overexpression but are tau aggregation dependent phenomena. Rapid nuclear targeting aligns with recent work demonstrating tau oligomers possess high conformational mobility and a tendency to interact with off-target proteins and hydrophobic domains such as those seen in the double layer nuclear envelope ^14,56,57^. These findings expand on previous observations by our lab that tau oligomers directly bind Lamin B2 and LBR ^20^, emphasizing that tau’s localization to nuclear boundaries and interaction with lamina components are not incidental but mechanistically integral to its neurotoxicity. Most importantly, the early emergence of tau oligomers ansd nuclear localization are consistent with theories affirming oTau as the primary toxic drivers in AD, preceding and potentiating neurofibrillary tangles ^11,13^. Our study supports the ongoing paradigm shift in AD research away from late stage accumulated fibrillary tau and amyloid beta plaques to early-stage oligomer mediated disruption ^58^.

Findings in PS19 tau transgenic mice paralleled those seen in cell culture. Nuclear invagination and pTau217 intensity were significantly more prevalent in aged PS19 mice, with trends in 5m PS19 resembling 9m WT, indicative of accelerated cellular aging seen in HGPS laminopathy and natural CNS neuron senescence ^47,48,59^. Furthermore, strong positive correlations between nuclear tau and lamina disruption only among PS19 mice affirm the nuclear toxic effects occurring under tau conditions. Perhaps the most profound consequence of lamina dysfunction is our observation of chromatin reorganization in PS19 mouse neurons. The nuclear lamina is critical in chromatin structure and stability via lamina-associated domains (LADs) which tether heterochromatin to the nuclear periphery ^49,52^. Our results indicate mouse neurons presenting with elevated tau-induced lamina intrusion show increased chromatin coverage and decreased heterochromatin density, particularly in the LAD rich NMZ. Disruption of LADs has been shown to correlate with altered transcriptional profiles and epigenome shifts characteristic of neuron dysfunction in AD ^50,51,54,60^. The observed chromatin decompaction in PS19 mice, a nuclear stress response, precede cell death and therefore are likely not secondary to neurodegeneration, again emphasizing that nuclear and chromatin disruption are early and causative.

Our most clinically relevant findings are from examination of human postmortem AD brain tissue. DAB staining finds concordance between emergence of MC1-positive misfolded tau and loss of LBR nuclear integrity beginning in Braak stage 3, well before large scale tau tangle deposition. The temporal alignment emphasizes that nuclear envelope disruption is precursor to late-stage degeneration and is critical in identifying nuclear lamina as a potential biomarker and therapeutic target in preclinical AD. Therapies aimed at lamina stabilization could mitigate chromatin and transcriptional dysregulation before degeneration takes root. Similar approaches hold precedent in progeria research where lamina stabilizing compounds have shown promise in mitigating nuclear pathology ^61^, but similar strategies adapted to AD tau induced nuclear aging warrants further research and exploration.

Perhaps most significantly is the broader similarities and applicability of the nuclear stress model to other neurodegenerative conditions. Nuclear envelope deformation and dysfunction are not confined to AD, as similar abnormalities are documented in Huntington’s disease where pathological huntingtin protein disrupts nuclear pore complexes and lamina structure ^30,62^. Certain other degenerative tauopathies including FTD and CTE also present with tau induced nuclear invagination and compromised nucleocytoplasmic trafficking ^18,26^. Collectively, these recent additions to the field point to a shared patho-mechanism of neurodegenerative laminopathy where diverse protein aggregates target nuclear vulnerability. Although fundamental precursors differ substantially, our study draws parallels between tau-induced lamina changes with farnesylation promoted laminopathy in HGPS ^49,59^. Nuclear phenotypes in early PS19 tau transgenic mice affirming premature emergence of nuclear invagination and chromatin loosening comparable to aged WT mirror the features of accelerated cellular aging.

In conclusion, our findings reframe some understanding of tau-mediated neurodegeneration by identifying the nuclear envelope as a substantial target of early tau toxicity. Our nuclear-centric model, supported by findings in human, mouse, and cell culture systems integrates structural biology, chromatin dynamics, and previous neurodegeneration understanding into a unified pathogenic cascade. Under this model, AD and potentially other neurodegenerative conditions are conceptualized as “nuclear envelope diseases,” involving aggregate-induced mechanical destabilization of genome integrity, thereby precipitating neuronal decline. By detailing these findings, this work served as compelling rationale for nuclear targeted therapeutic strategies aimed at restoring lamina function and genome integrity. Our findings corroborate and extend existing literature on mechanical perspectives of AD research and redirects attention toward previously underappreciated nuclear targets.

## LIMITATIONS AND FUTURE DIRECTIONS

While our findings offer considerable mechanistic insight, several limitations warrant acknowledgement. First, the optogenetic system is a powerful tool for temporally controlling and observing tau oligomerization, but it may not fully recapitulate the complex environment of tau accumulation in vivo. This problem is fairly mitigated through use of additional AD models including transgenic mice and human AD postmortem brain tissue. Next, our primary focus of nuclear disruption centered around the protein targets LB2 and LBR, and additional components of the nuclear envelope, a highly complex and dynamic cell structure, merit further investigation, including LINC proteins Emerin and SUN1/2, as well as NPC components. Current work using co-immunoprecipitation to resolve precise molecular interactions between tau and LB2, LBR, and other components is underway. Furthermore, chromatin decompaction was observed, but future work using single-cell RNA sequencing and proteomics profiling is aimed at mapping temporal relationships between lamina disruption, chromatin alterations and transcriptional deregulation. Finally, protein tracking by Turbo-ID proximity labeling will be used to monitor changes in nucleocytoplasmic trafficking as a result of nuclear disruption. Ultimately, further investigation into events linking tau oligomerization, nuclear dysfunction, and downstream cellular degeneration will provide deeper insight into developing effective therapeutic interventions and preventing neurodegeneration in AD.

## MATERIALS AND METHODS

### Compliance statement

All research was performed with approval from University of Virginia School of Medicine. The use of animals was approved by the University of Virginia Institutional and Animal Care and Use Committee (PUBLIC HEALTH SERVICE (PHS) ASSURANCE #A3245-01; USDA REGISTRATION #: 52-R-0011; Protocol#: 4454). All animals were housed in IACUC-approved vivariums at UVA School of Medicine.

### Cell culture

All cell cultures were maintained at 37 °C with 5% CO2. All cell counts were performed in duplicate using the RWD Cellometer with 0.4% trypan blue dye (Thermo Fisher).

iPSC neuronal cell differentiation and maturation: Human iPSC were obtained from JAX iPSC Collection (Catalog#: JIPSC001000). Human iPSC derived neural progenitor cells (NPCs) were differentiated with STEMdiff™ Forebrain Neuron Differentiation Media (Stem Cell Tech cat#08600) on Corning® Matrigel® coated tissue culture treated plates. NPCs were passaged and plated at 50,000 cells/cm2 in with the maintained in serum-free STEMdiff™ Neural Progenitor Medium 2 (Stem Cell Tech cat#08560) on Corning® Matrigel® hESC-qualified Matrix (Corning cat#354277) coated tissue culture plates. NPCs were plated at 50,000 cells/cm2 and passaged at 90% confluency by AccutaseTM (Stem Cell Tech cat#07920) dissociation as necessary. A full media change was performed every other day. Low passage (passage < 3) NPCs were cryopreserved in STEMdiff™ Neural Progenitor Medium 2 with 10% DMSO, and all NPCs used for experimentation were maintained at passage <6. For the maturation of neurons, the NPCs were transduced with a NEUROG2 lentivirus (GeneCopoeia cat#LPP-T7381-Lv105-A00-S) at MOI 3 to induce iPSC-derived neuronal cells (hiNC). After 24 h of transduction, a full media change was performed.

### Human iPSC induced neurons cell culture and OptoTau lentivirus transduction

Prepare four 75 cm2 flasks, designated as mCherry 0 min, mCherry 60 min, OptoTau 0 min, and OptoTau 60 min. Each flask should contain 8 mL of DMEM/F-12 medium (STEMdiffTM, #36254, USA) supplemented with 40 µL Matrigel (Corning®, #354277, USA) and be pre-coated for 2 hours before cell passaging. After pretreatment, dissociate and passage the cells, resuspending them in Neural Progenitor Medium (STEMdiffTM, #05834, USA) at a density of 6×10⁵ cells/mL, then transfer the suspension to the coated flasks. Upon cell adhesion, initiate lentiviral transduction by adding 4 µL/mL mCherry lentivirus to the mCherry groups or 4 µL/mL OptoTau lentivirus to the OptoTau groups. After 48 hours, repeat the lentiviral treatment and include 1 µL/mL NEUROG2 lentivirus particles (GeneCopoeia, CLP-HPRM50462, USA), simultaneously replacing the medium with Forebrain Neuron Maturation Medium (STEMdiffTM, #08605, USA). Maintain the cultures by refreshing the medium every three days. On Day 21, subject the mCherry 60 min and OptoTau 60 min groups to 488 γ blue light irradiation for 60 minutes, then harvest the cells for lysate collection.

### Live cell staining with NucSpot Live 488

Repeat human iPSC derived neuron cell culture and OptoTau transduction using 35mm glass bottom petri dish without prolonged blue light exposure. 1 hour prior to live cell imaging of induced neurons, cells were stained with NucSpot Live 488 (Biotium Cat #40081), a low toxicity DNA nuclear stain designed for live cell imaging. Cells were incubated in Nucspot 1000x diluted to 1x concentration in neuronal forebrain maturation medium for 15-20 min at 37°C. Dilution was supplemented with 25 μM Verapamil to slow nuclear dye leakage. Following incubation, dilution was removed and replaced with regular forebrain maturation medium and incubated for 20-30 minutes to allow optimal pH under 5% CO_2_ to be reached. To maintain pH during live cell imaging experiment, medium was supplemented with 25 mM HEPES buffer solution following incubation.

### Immunocytochemistry (ICC) staining of fixed iPSC derived neurons

In glass bottom 24 well plates, cells were fixed with 0.5 ml 4% PFA/PBS for 10 minutes and washed 3x in PBS for 5 min. Cells were permeabilized in 0.5ml PBS/0.1% Triton X-100 (PBST) for 15-30 min and blocked in 0.5ml of 5% BSA - 5% goat Serum in PBST for 1 hour. Cells were then incubated in 1° antibody diluted in 5% BSA/PBST overnight at 4°C. Plates were wrapped in Parafilm and photobleached by led lamp. The next day, the cells were washed 3x in PBS-T, 10 min each before being incubated in 2° antibody diluted 1:800 in 5% BSA/PBST, 2 hrs at RT. In all remaining steps, cells were covered in foil to prevent ambient light photobleaching. For antibody information and dilutions, view materials section. After 2° antibody, cells were washed 1x in PBS-T for 10 min and incubated in DAPI diluted 1:10,000 in PBST (5 mg/ml stock solution) for 15 min after first wash. Then cells were washed 2x with PBST, and once of PBS, 10 min each. 12 μl Prolong Gold Antifade mounting media was dropped in the middle of each well and covered with 12mm coverslips. Plates were covered from light and dried in the fume hood overnight before long term storage at 4°C.

### Immunohistochemistry (IHC) staining of fixed brain tissues with fluorescence

Brain tissue sections were selected from 40μm slices of fixed post-mortem human brain tissues and dissected mouse brain tissues stored at 4°C in 0.01% sodium Azide/PBS. Sections were transferred to 24 well plates at 1 section per well. Sections were first washed in PBS for 10 mins and permeabilized in 0.5ml PBS/0.25% Triton X-100 (PBST) for 15-30 min. Sections were blocked in PBS-T with 5% BSA and 5% Goat Serum for 1h at room temperature (RT) and transferred to 1° antibody diluted in 5% BSA/PBST and wrapped in parafilm before incubation overnight at 4°C. During incubation, sections were photobleached by LED lamp. The next day, sections were washed 3x in PBST (10 min each) and incubated in 2° antibody diluted 1:800 in 5% BSA/PBST for 2 hrs at RT. For antibody information and dilutions, view materials section. In remaining steps, plates are covered in foil to prevent ambient photobleaching. After 2°, sections are washed 1x in PBS-T for 10 min and incubated in DAPI diluted 1:5,000 in PBST (5 mg/ml stock solution) for 15 min after first wash. Sections are then washed 2x with PBST and washed with PBS upon mounting to glass slides. 50 μl of Prolong Gold Antifade mounting media was dropped on each slide and covered with 12mm glass cover slips. Slides were covered placed in dark slide box died overnight at RT in the fume hood before long term storage at 4°C.

### Immunohistochemistry single stain of fixed tissues with DAB (3,3’-diaminobenzidine)

Brain tissue sections were selected from 40μm slices of fixed post-mortem human brain tissues and dissected mouse brain tissues stored at 4°C in 0.01% sodium Azide/PBS. Sections were washed in PBS 1x for 10 minutes and treated with 1% H_2_O_2_ diluted in dH_2_O for 15 mins to inactivate endogenous peroxidase activity. Sections were then rinsed 2x in PBS, 15 min each, and blocked in 0.4% triton X-100/PBS, 1% Bovine Serum Albumin (BSA), 4% Normal Goat Serum (NGS) for 30 min. Sections were then transferred to 1° antibody diluted in DAKO antibody diluent (5% BSA in 0.25% TritonX-100/PBS) and incubated overnight at 4°C with photobleaching. For antibody information and dilutions, view materials section. The next day, tissue sections were rinsed 2x in PBS for 15 min each. Sections were then incubated for 1h at RT in BTA solution (0.44% Biotinylated goat anti-rabbit / mouse (depending on 1° antibody) IgG diluted in 0.3% triton X-100/PBS) and washed 2x in PBS for 15 min each. Sections were treated with Avidin Biotin Complex (ABC) (0.88% Avidin, 0.88% Biotin diluted in 0.3% triton X-100/PBS, prepared 30 minutes prior to use). Following ABC incubation, sections were washed 3x in PBS, 10 min each wash. DAB solution was prepared while washing with 1 tablet DAB, 1 tablet Urea (Sigma-Aldrich Cat# D4193) diluted in 5 mL dH_2_O and vortexed until fully dissolved. Sections were treated in DAB solution one at a time for 1-3 min until section turned medium brown color and immediately placed in PBS. Sections were then washed 2x in PBS, 5 min each wash and mounted to glass slides. Slides were dried overnight at 37° C. The following day, slides were dehydrated by washing in 70% EtOH, 85% EtOH, 95% EtOH, and 100% EtOH for 2 min each followed by 100% EtOH for 5 min. Slides were then cleared by washing 2x in Xylene, 5 min each, followed by immediately dropping 50 μl Permout Mounting Medium (Fisher Scientific Cat# SP15-500) and covering with glass coverslips. Slides were dried overnight in the fume hood at RT and stored at 4°C for long term storage.

### Primary Antibody Information

| Name | Anti | Target | Dilution Factor | Storage | Source | Catalogue |
| --- | --- | --- | --- | --- | --- | --- |
| LBR Rabbit PolyAb | Rabbit |  | 1:500 IHC;<br>1:1000 ICC | -20°C | Proteintech | 12398-1-AP |
| AT8 Phosphotau Monoclonal Antibody | Mouse | Phosphorylated tau | 1:1000 WB;<br>1:500 IHC;<br>1:500 ICC | -20°C | Invitrogen | MN1020 |
| GAPDH<br>Mouse<br>McAb | Mouse | GAPDH<br>(Control) | 1:1000 WB | -20°C | Proteintech | 60004-1-Ig |
| Lamin B2<br>Rabbit mAb | Rabbit | LaminB2 | 1:1000 WB | -20°C | Cell<br>Signaling<br>Tech | E1S1Q |
| Lamin B2 | Mouse | LaminB2 | 1:400 IHC;<br>1:500 ICC | -20°C | Invitrogen | 33-2100 |
| Tau5 | Mouse | Phosphorylated<br>tau | 1:400 IHC;<br>1:400 ICC | -20°C | Millipore | MAB361 |
| PTau 217<br>Polyclonal<br>Ab | Rabbit | Phosphorylated<br>tau | 1:400 IHC;<br>1:500 ICC | -20°C | Invitrogen | 44-744 |
| TOMA2 | Mouse | Phosphorylated<br>oligomeric tau | 1:500 IHC | -20°C | Collaborator |  |
| Tuj1 Beta<br>Tubulin III | Chicken | Neurons | 1:500 ICC | 4°C | Aves | TUJ |
| MAP-2 | Chicken | Neuronal<br>processes | 1:400 IHC | 4°C | Aves | MAP |
| mCherry<br>mAb (16D7) | Rat | mCherry | 1:500 ICC | -20°C | Invitrogen | M11217 |
| Dapi* | Stain | Nuclei | 1:5000 IHC;<br>1:10000 ICC | -20°C | Collaborator |  |
| CP13 | Mouse | anti-tau p202<br>(early) | 1:500 WB;<br>1:300 IHC | -20°C, -<br>80°C | Collaborator |  |
| MC1 | Mouse | misfolded tau | 1:500 WB;<br>1:300 IHC | -20°C, -<br>80°C | Collaborator |  |
| PHF-1 | Mouse | anti-tau pS396<br>(late); insoluble<br>paired helical<br>filaments<br>(PHFs) | 1:500 WB;<br>1:300 IHC | -20°C, -<br>80°C | Collaborator |  |

### Secondary Antibody Information

| Name | Animal | Type | Dilution Factor | Storage | Source | Catalogue |
| --- | --- | --- | --- | --- | --- | --- |
| Goat anti-Rabbit IgG<br>(H+L) | Rabbit | Alexa Fluor 488 | 2mg/mL | 4°C | Invitrogen | A11008 |
| Goat anti-Rabbit IgG<br>(H+L) | Rabbit | Alexa Fluor 594 | 2mg/mL | 4°C | Invitrogen | A11012 |
| Goat anti-Rabbit IgG (H+L) | Rabbit | hRPC | 1:5000 WB | -20°C | Invitrogen | G21234 |
| Goat anti-rat IgG (H+L) | Rat | Alexa Fluor 488 | 2mg/mL | 4°C | Invitrogen | A11006 |
| Goat anti-Rat IgG (H+L) | Rat | Alexa Fluor 594 | 2mg/mL | 4°C | Invitrogen | A11008 |
| Goat anti-rat IgG (H+L) | Rat | hRPC | 1:5000 WB | -20°C | Invitrogen | A10549 |
| Goat anti-Mouse IgG (H+L) | Mouse | Alexa Fluor 405 | 2mg/mL | 4°C | Invitrogen | A48255 |
| Goat anti-Mouse IgG (H+L) | Mouse | Alexa Fluor 488 | 1mg/mL | 4°C | Invitrogen | A28175 |
| Goat anti-Mouse IgG (H+L) | Mouse | Alexa Fluor 594 | 2mg/mL | 4°C | Invitrogen | A11032 |
| Goat anti-Mouse IgG (H+L) | Mouse | hRPC | 1:5000 WB | -20°C | Invitrogen | G21040 |
| Goat anti-Chicken IgG (H+L) | Chicken | Alexa Fluor 488 | 2mg/mL | 4°C | Invitrogen | A11039 |
| Goat anti-Chicken IgG (H+L) | Chicken | Alexa Fluor 594 | 2mg/mL | 4°C | Invitrogen | A11007 |
| Goat anti-Chicken IgG (H+L) | Chicken | Alexa Fluor Plus 647 | 2mg/mL | 4°C | Invitrogen | A32933 |
| Goat anti-Chicken IgY (H+L) | Chicken | hRPC | 1:5000 WB | -20°C | Invitrogen | A16054 |

### Quantification and statistical analysis

#### Image analysis

Fluorescence labeling intensity was measured using FIJI (ImageJ). Colocalization of tau isoforms and lamina components was analyzed using Coloc2 plugin for FIJI. Quantification of nuclear invagination rates by ROI was done using a custom plugin ‘Nuclear Laminin Distribution’ for FIJI provided by Paonessa F. at the Livesey Lab ^25^. Plugin uses dsDNA dapi staining channel to map nuclear bounds and lamina channel to map internal and peripheral lamin components. Nuclei with an internal lamina area > 0.3 of total lamin are counted as invaginated. Additional channels can be used to measure mean intensity within the nuclear bounds. For EM quantification, invagination size was measured by relative membrane length using FIJI. Relative chromatin coverage was measured using FIJI. Chromatin clump density was measured using the ‘connected components labeling’ tool from the MorpholibJ plugin for FIJI ^63^. Clumps were quantified as closely pixels with an area ≥ 0.04 μm^2^ and normalized to nucleus area.

#### Statistical analysis

Statistical analyses and figures artwork were performed using GraphPad Prism version 10.00 for Windows with a two-sided α of 0.05. All group data are expressed as mean ± SEM. Column means were compared using one-way ANOVA with treatment as the independent variable. Group means were compared using two-way ANOVA using factors of genotype and fraction treatment, respectively. When ANOVA showed a significant difference, pair wise comparisons between group means were examined by Tukey’s, Dunnett or uncorrected Fisher’s LSD multiple comparison test. Significance was defined when p < 0.05.

## Acknowledgments

Brain tissues were provided by the Emory University Goizueta Alzheimer’s Disease Research Center (P30 AG066511). We would like to thank the following funding agencies for their support: to L.J., NIH/NIA (R01AG091577), UVA Provost Award, UVA Health System, UVA Brain Institute Pitch and Catch Fund, Jim and Bruce Eck Fund, Strang Neuroscience Research Award, American Federation for Aging Research.

## Author contributions

Conceptualization: L.J. and A.E.

Methodology: N.E., S.Y., R.R., E.S., W.G., Q.W., A.E., L.J.

Investigation: N.E., S.Y., R.R., E.S., W.G., Q.W., A.E., L.J.

Visualization, N.E., S.Y., R.R., E.S., Q.W., A.E., L.J.

Writing: N.E., S.Y. and L.J.;

Editing: L.J., N.E., S.Y., R.R., E.S., W.G., Q.W., A.E.;

Supervision: L.J.

Funding Acquisition: L.J.

## Declaration of interests

The authors declare no competing interests.

## Data and code availability

All data reported in this paper will be shared by the lead contact upon request.

## Notes

### Competing Interest Statement

The authors have declared no competing interest.

